# Sex-stratified fixel-based analysis reveals reduced frontal fibre density in males with first-episode of psychosis

**DOI:** 10.64898/2026.09.15.751722

**Authors:** Stella M. Sánchez, Helmut Schmidt, Antonín Škoch, Kenneth Hugdahl, Filip Španiel, Jaroslav Hlinka

**Author notes:** Corresponding author: Stella M. Sánchez, Pod Vodárenskou věží 2, 182 00 Praha 8, Czech Republic.

## Abstract

**Background:** Schizophrenia has been conceptualised as a disorder of brain disconnectivity, with white matter abnormalities contributing to altered neural communication. However, fibre-specific white matter alterations in early illness and their potential sex-related differences remain insufficiently characterised. We applied fixel-based analysis to investigate sex-stratified white matter microstructural alterations in first-episode schizophrenia (FES).

**Methods:** Diffusion MRI data from individuals with FES and healthy controls were stratified by biological sex. The male cohort included 79 patients and 29 controls, and the female cohort included 48 patients and 46 controls. Fixel-based analysis was performed separately in males and females to assess fibre density, fibre-bundle cross-section, and their combined measure. Head motion was included as a covariate in the fixel-based analyses. A descriptive follow-up voxel-scale analysis extracted mean fibre density from a binary mask derived from significant male fixels and applied the same region to the female cohort.

**Results:** Males with first-episode schizophrenia showed significantly reduced fibre density compared with male controls, primarily in bilateral frontal white matter and a thin segment of the anterior corpus callosum. The voxel-scale follow-up confirmed lower mean fibre density in males within the significant fixel-derived mask. In contrast, females showed no significant group differences in any fixel-based metric or within the male-derived region.

**Conclusions:** Early white matter microstructural alterations were detected in males with FES but not in females. These findings highlight a differential pattern emerging from sex-stratified analyses and support further investigation of sex-related differences in white matter alterations during the early stages of schizophrenia.

## 1 Introduction

Schizophrenia is a severe and chronic mental disorder characterised by recurrent psychotic episodes, which affects both men and women. Accumulating evidence suggests that biological sex influences how individuals experience the disorder; for example, men and women differ in risk factors, age at onset, prevalence, symptom severity, and treatment outcomes (1–3). However, the existence and extent of true sex- or gender-related differences in schizophrenia remain debated, partly due to methodological heterogeneity and confounding factors (e.g., medication exposure, illness chronicity, and sampling differences) (2,4). A better understanding of how sex and gender influence the neurobiology of schizophrenia may contribute to improved disease management and the development of more targeted therapeutic strategies. Here, we use “sex” to refer to biological sex as recorded at recruitment.

One of the most widely supported theoretical frameworks conceptualises schizophrenia as a disorder of brain disconnectivity, involving altered or inefficient integration between distributed brain regions. This framework implicates not only grey matter abnormalities but also disruptions in white matter (WM) pathways that support long-range neural communication (5–7). WM abnormalities, reflecting altered tissue organisation and microstructure, can be investigated using diffusion-weighted imaging (DWI), a non-invasive in vivo neuroimaging technique sensitive to the diffusion of water molecules and capable of inferring tissue architecture. Most DWI studies rely on voxel-derived diffusion metrics (8,9), such as fractional anisotropy (FA) and mean diffusivity (MD), to characterise WM microstructure.

While informative, these voxel-wise measures are inherently descriptive and cannot disentangle multiple fibre populations within a single voxel (10). Estimates suggest that 30–90% of WM voxels contain crossing fibres (11); therefore, this limitation can obscure biologically meaningful alterations. To overcome this constraint, Fixel-Based Analysis (FBA) (12) was developed as a statistical framework that enables the modelling and quantification of fibre-specific WM properties within individual voxels. FBA introduces the concept of a fixel, defined as a distinct fibre population within a voxel, thereby providing a more anatomically precise and biologically interpretable characterisation of WM organisation. Accordingly, FBA offers greater anatomical specificity than voxel-averaged metrics.

Previous DWI studies have reported WM microstructural alterations across different stages of schizophrenia, including both chronic patients and individuals experiencing a first episode of schizophrenia (FES). Compared with healthy controls, individuals with schizophrenia generally exhibit lower FA in the prefrontal and temporal lobes and in the WM tracts connecting these regions (6), as well as in the splenium of the corpus callosum (13) and the cingulum bundle (14). Additionally, higher global MD values have been reported in schizophrenia patients (13). Findings in FES are less consistent across studies (15,16), potentially reflecting early-stage neurobiology, sample differences, and treatment effects. Some studies have reported higher FA and lower MD in regions such as the right superior longitudinal fasciculus and the corpus callosum compared with healthy controls (14), which may reflect early or compensatory microstructural changes. Despite extensive voxel-wise DWI work, fibre-specific alterations—particularly in early illness—remain less well characterised.

In the present study, we conducted sex-stratified analyses to examine WM microstructure in individuals with first-episode schizophrenia, an illness stage that may minimise confounding effects related to chronicity and prolonged medication exposure. The male cohort consisted of 79 patients with FES and 29 healthy male controls, while the female cohort included 48 patients with FES and 46 healthy female controls, all recruited during the same period. Using FBA, we aimed to identify fibre-specific WM abnormalities associated with the early stage of illness and to assess whether these alterations differ by sex. To our knowledge, this is the first sex-stratified FBA study in FES to investigate WM alterations at the fixel level, providing novel insights into sex-specific patterns of WM pathology in schizophrenia.

## 2 Methods and Materials

### 2.1 Participants

All participants were recruited as part of the Early-Stage Schizophrenia Outcome (ESO) study, a large ongoing prospective programme that investigates individuals experiencing a first episode of schizophrenia spectrum disorders and is coordinated by the National Institute of Mental Health (NIMH) based in the Czech Republic. The study was approved by the Ethics Committee of the Prague Psychiatric Centre on June 29, 2011 (protocol code 69/11) and was conducted in accordance with the most recent version of the Declaration of Helsinki. All participants received a comprehensive explanation of the study procedures and provided written informed consent before enrollment. To ensure privacy and confidentiality, all sensitive participant data were anonymised.

For this study, we analysed data from the baseline visit that occurred at the first episode of psychosis. The diagnosis was confirmed by two experienced psychiatrists based on the ICD-10 Diagnostic Criteria for Research (WHO, 1992). Inclusion criteria required that participants: (1) were undergoing their first psychiatric hospitalisation; (2) had an ICD-10 diagnosis of schizophrenia, acute and transient psychotic disorders, or schizoaffective disorders, as determined by the Mini-International Neuropsychiatric Interview (MINI (17)); (3) had experienced fewer than 24 months of untreated psychosis; and (4) were at least 18 years old. Individuals diagnosed with psychotic mood disorders, including bipolar disorder or major depressive disorder with psychotic features, were excluded from the study.

In addition, a healthy control (HC) group was recruited through local advertisements, and 81 individuals were included in the study. The main exclusion criteria for HC were a personal history of any psychiatric disorder or substance abuse, as assessed using the MINI, and a family history of psychotic disorders in first- or second-degree relatives. Additional exclusion criteria for both groups included current neurological disorders, a history of seizures or head injury with altered consciousness, intracranial haemorrhage or neurological sequelae, intellectual disability (IQ < 80), history of substance dependence, and any contraindication for MRI scanning (18).

All participants in this study were recruited and scanned between March 2016 and January 2021. The MRI protocol was identical for all individuals and was conducted at the same MRI facility (NIMH in Klecany, Czech Republic). All FES individuals were treated with antipsychotics since their hospitalisation and underwent MRI scanning during the first month at the hospital. A total of 132 individuals with a first episode of schizophrenia (FES) and 81 HC subjects were included in this study, considering both men and women. After a visual inspection of the data and the sex-stratification, we obtained the following groups (with final group size n): male-FES (n=79), male-HC (n=29), female-FES (n=48), and female-HC (n=46).

### 2.2 MRI Acquisition and Preprocessing

MRI data were acquired using a 3T Siemens Prisma scanner.

DWI parameters were: spin-echo EPI; TR/TE = 8300/84 ms; voxel size 2 mm isotropic; flip angle 90°; six b0 volumes in each phase-encoding direction; two shells (b = 1000, 2500 s/mm^2^) with 64 directions each. T1-weighted sequence was: MPRAGE: TR/TE/TI = 2300/4.63/900 ms; flip angle 10°; voxel size 1 mm isotropic; FOV 256 mm; GRAPPA 2.

All data modalities underwent visual quality control. Toolboxes and commands from MRtrix3 and FSL software packages were used. DWI preprocessing consisted of denoising, Gibbs ringing removal, susceptibility correction (TOPUP), eddy-current/head-motion correction (EDDY), and intensity inhomogeneity correction (N4, ANTs). WM and CSF intensities were normalised across participants. Fibre orientation distributions (FODs) were estimated using the multi-shell, multi-tissue (MSMT) Constrained Spherical Deconvolution (CSD) method (19).

On the other hand, T1w images were coregistered to DWI using boundary-based registration, skull-stripped with BET, and segmented using FAST (FSL). More details about the preprocessing can be found in (20).

### 2.3 Fixel-based Analysis (FBA)

FBA enables the assessment of fibre-specific WM properties through three metrics: fibre density (FD), related to the intra-axonal restricted compartment at each fixel; fibre-bundle cross-section, sensitive to macrostructural and morphological changes; and finally, a combined measure of fibre density and cross-section (21).

In this study, FBA was applied to assess WM microstructural alterations in FES in both sexes separately (i.e. male-FES vs male-HC, and female-FES vs female-HC). Connectivity-based fixel enhancement (CFE) was performed using 4 million streamlines from a 20-million-streamline tractogram template filtered by spherical-deconvolution-informed filtering of tractograms algorithm. All CFE parameters were set to default use (10 mm FWHM smoothing; C=0.5; E=2; H=3), and non-parametric permutation testing (5000 permutations) produced family-wise error–corrected *p*-values. In addition, a covariate column was added to the general linear model to control for the effect of the absolute head motion of the participants. This confounding variable is defined as the RMS displacement from the first volume, where large values (>2–3 mm) usually indicate a restless participant (22), and was obtained from EDDY correction during preprocessing.

To assess the empirical false-positive rate of our FBA pipeline, we performed a null-validation resampling control within the male cohort. We randomly partitioned the full male sample into two groups and repeated the full FBA workflow 100 times. An iteration was considered “positive” if any significant effect was detected in FD. Across 100 random splits, 5 iterations yielded at least one significant finding, consistent with the nominal 5% Type I error rate expected under the null hypothesis and suggesting no systematic inflation of false positives in our FBA pipeline.

### 2.4 from Fixel to Voxel Scale

Although voxel-based approaches have important limitations when used as the primary framework for analysing WM organisation, representing the results at the voxel scale may offer some practical advantages. In particular, voxel-wise maps can be more easily subjected to linear and non-linear spatial transformations, without the additional complexity of transforming fibre orientations encoded at the fixel level. This makes voxel-scale representations useful, for example, when comparing results obtained in different template spaces.

As a follow-up analysis to the FBA, we evaluated whether the fixel-based effects observed in the male cohort were also detectable when represented at the voxel scale. To this end, we first built a binary mask from the significant FBA results in the male cohort. Voxels were assigned a value of 1 if they contained at least one significant fixel (*p* < 0.05), and a value of 0 otherwise. This mask, therefore, represented the spatial extent of the significant fixel-based findings at the voxel level.

For each participant, fixel-wise FD values were converted into a voxel-wise FD image by averaging FD values across all fixels within each voxel. The resulting voxel-wise FD image was then multiplied by the binary mask, yielding an FD map restricted to the region identified by the significant FBA findings. From these participant-specific masked FD maps, we extracted the mean FD value within the mask for subsequent statistical analyses, as described in the next section.

Finally, the same male-derived binary mask was applied to the female cohort to assess whether the WM region showing significant effects in males also showed evidence of group differences in females. The same fixel-to-voxel conversion and masking procedure was applied to each female participant, and mean FD values within the mask were compared between the HC and FES female groups.

### 2.5 Statistical Analysis

For demographic and MRI-derived measures, normality and homogeneity of variances were assessed using the Kolmogorov–Smirnov test and a two-sample *F*-test, respectively. When assumptions were met, group differences were evaluated using Student’s *t*-tests; otherwise, the non-parametric Mann–Whitney U test was applied. Since these comparisons were intended to identify potential confounders rather than to test primary hypotheses, *p*-values for demographic and head-motion variables were reported uncorrected; variables showing at least an uncorrected significant between-group difference were considered for inclusion as covariates in the subsequent neuroimaging analyses. All statistical analyses were performed using MATLAB (MathWorks, Natick, MA, USA).

For the voxel-scale microstructural follow-up analysis, group differences in mean FD values extracted from the male-derived binary mask were assessed separately within each cohort. Specifically, mean FD within the mask was compared between HC and FES males, and then between HC and FES females, using Student’s t-tests when assumptions of normality and homogeneity of variances were met, and Mann–Whitney U tests otherwise. No covariates were included in these voxel-scale follow-up comparisons. Effect sizes were quantified using Cohen’s d.

To further evaluate how well FD within the male-derived binary mask discriminated between groups within each cohort, Receiver Operating Characteristic (ROC) curves were computed using the mean FD value extracted from the mask for each participant. For each ROC curve, we calculated the true positive rate (TPR), corresponding to *sensitivity*, and the false positive rate (FPR), defined as *1 − specificity*. Discriminative performance was quantified using the area under the ROC curve (AUC).

In addition, we conducted a sensitivity power analysis to aid the interpretation of the non-significant findings in the female cohort. Power was evaluated for an independent two-sample t-test, two-sided, α = 0.05, using the observed female sample sizes (HC: *n*=46; FES: *n*=48). Specifically, we quantified the probability of detecting a group difference in females across a range of assumed standardised effect sizes (Cohen’s *d*), focusing on three scenarios: (i) an effect matching that observed in males within the male-defined binary mask (*d*=1.3), (ii) a conservative scenario corresponding to half the male effect (*d*=0.65) to partially mitigate potential inflation of the male effect due to mask selection, and (iii) a moderate effect size (*d*=0.5). Power estimates were computed in MATLAB using the noncentral-*t* framework. In addition, we derived the effect size required to achieve 80% power (*d*_*80%*_) by computing power over a dense grid of *d* values and interpolating the value at 0.80.

## 3 Results

### 3.1 Demographic and MRI Data

Table 1 summarises the statistical analysis of demographic and MRI-related variables. No significant between-group differences in age were observed in either the male or female cohorts. Years of education (YoE) differed significantly between FES and HC in the male cohort (*p* = 1.17 × 10^-4^), but not in the female cohort (*p* = 0.083). The lower educational attainment observed in males with FES is consistent with previous reports showing fewer years of education in individuals with schizophrenia compared with healthy controls (23). Absolute head motion (AbsMot) also differed significantly between groups in the male cohort (*p* = 0.039), whereas no significant difference was observed in the female cohort (*p* = 0.572). Because head motion can introduce systematic artefacts in diffusion MRI data and potentially bias between-group comparisons (24), AbsMot was included as a covariate in the subsequent analyses. In contrast, YoE was not included as a covariate, as reduced educational attainment is commonly reported in schizophrenia (23,25) and may reflect disorder-related characteristics rather than an independent confounding factor.

**Table 1:** Demographic and MRI Data of HC and FES Participants.

| Metric/Cohort | HC | FES | $p$ value |
| --- | --- | --- | --- |
| Male Cohort | n = 29 | n = 79 |  |
| Age (yrs) | 30.15 +/- 6.58 | 27.29 +/- 6.87 | 0.0558 |
| YoE (yrs) | 17.13 +/- 2.65 | 14.49 +/- 3.17 | $1.17 \times 10^{-4}$ |
| AbsMot (mm) | 0.717 +/- 0.515 | 0.99 +/- 0.77 | 0.039 |
| Female Cohort | n = 46 | n = 48 |  |
| Age (yrs) | 30.27 +/- 10.03 | 28.67 +/- 7.18 | 0.371 |
| YoE (yrs) | 16.0 +/- 2.79 | 14.9 +/- 3.12 | 0.083 |
| AbsMot (mm) | 0.83 +/- 0.51 | 0.90 +/- 0.81 | 0.572 |

*Abbreviations: HC, Healthy Controls; FES, First-episode Schizophrenia; YoE, Years of Education; AbsMot, Absolute Motion. All $p$ -values reported are before multiple corrections.*

In addition, no significant differences between males and females were observed in age at onset or clinical symptom severity. Specifically, PANSS Positive, Negative, and General scores were comparable between sexes, as was age of symptoms onset (all *p* > 0.05; Table 2). These findings indicate that the male and female patient groups did not significantly differ in these clinical characteristics.

**Table 2:** Clinical Data of FES Participants. Comparisons between Male and Female Cohorts.

| Metric | Male<br>(mean ± SD) | Female<br>(mean ± SD) | *p* value | Test statistic |
| --- | --- | --- | --- | --- |
| Age of S. onset<br>(years) | 26.32 ± 6.87 | 28.04 ± 7.25 | 0.224 | $U = 1651.0$ |
| PANSS Positive | 11.06 ± 3.48 | 10.98 ± 3.54 | 0.849 | $U = 1857.5$ |
| PANSS Negative | 16.24 ± 5.73 | 15.52 ± 5.90 | 0.574 | $U = 1782.5$ |
| PANSS General | 27.89 ± 6.89 | 28.85 ± 7.39 | 0.457 | $t = -0.747$ |

Abbreviations: PANSS, Positive and Negative Syndrome Scale. All *p*-values reported are before multiple corrections.
Values are presented as mean ± standard deviation (SD). Between-sex differences were assessed using the Mann–Whitney *U* test for age at onset, PANSS Positive, and PANSS Negative scores, and an independent-samples *t*-test for PANSS General scores. *U* denotes the Mann–Whitney test statistic and *t* the Student's *t*-test statistic.

### 3.2 FBA on Male Cohort

The male-FES group showed significantly reduced fibre density (FD) compared with male healthy controls. These differences were primarily in fixels located in frontal WM regions of both hemispheres and included a thin segment of the anterior corpus callosum. All effects survived FWER correction, with *p*-values ranging from 0.03 to 0.05. Fixels corresponding to this *p*-value range will be referred to as ‘significant fixels’.

Figure 1A illustrates the streamlines corresponding to significant fixels, coloured by principal fibre orientation (RGB-XYZ convention). In Figure 1B, the same streamlines are displayed using a colour scale that represents the percentage FD reduction relative to the HC mean. Yellowish tones indicate regions where the FES group shows approximately 50% lower FD than HC.

**Figure 1.**
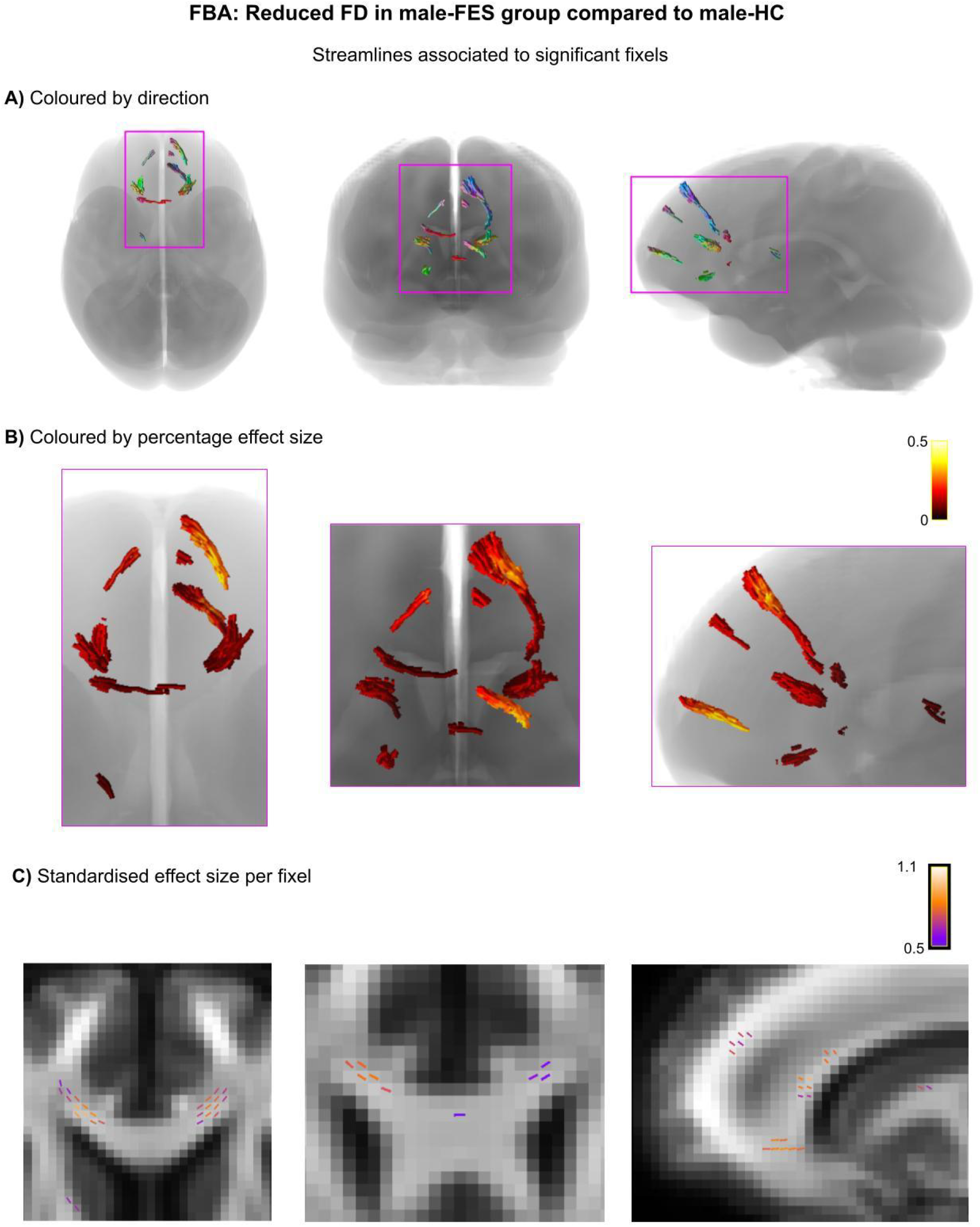
Outcomes of FBA in the male cohort showed reduced FD in FES compared to HC. **(A)** Streamlines corresponding to significant fixels, coloured by principal fibre orientation (RGB–XYZ convention), shown in axial, coronal, and sagittal views. (B) Zoomed-in views of the same region, with streamlines coloured according to the percentage FD reduction relative to the HC mean (hot colour scale). Yellowish colours indicate the percentage effect size ∼50%. (C) Representative axial, coronal, and sagittal cropped slices displaying significant fixels coloured by standardised effect size (Cohen’s d). Abbreviations: FBA, Fixel-Based Analysis; FD, Fibre Density; HC, Healthy Controls; FES, First-Episode Schizophrenia.

To characterise the magnitude of these effects, we extracted standardised effect sizes (Cohen’s *d*) for all significant fixels. As an example, Figure 1C displays the significant fixels from one slice in the three anatomical views: axial, coronal, and sagittal. Values ranged from 0.50 to 1.16, indicating moderate effects in purple fixels and strong group differences in light orange fixels. To simplify the visualisation, we selected a slice containing one significant fixel per voxel; however, it should be noted that, across the whole brain, each voxel may contain more than one fixel.

We further computed each participant’s mean FD across the binary mask defined from significant fixels (Figure 2A). As expected from the way the mask was defined, mean FD was lower in the FES group at the voxel scale (FD_HC_=1.10 ± 0.11; FD_FES_=0.97 ± 0.09). After verifying normality and homogeneity of variance, a two-sample *t*-test revealed a highly significant group difference (*p*=3.46 × 10^-8^, *d*=1.29); note that while the effect size is likely inflated by the region selection procedure, it provides a basic robustness validation that the signal is indeed visible at the voxel level and simplifies comparability across datasets. Figure 2B displays the distributions of mean FD values for each group (green for HC and pink for FES). This analysis was therefore intended to summarise and visualise the fixel-based effect at the voxel scale, rather than to provide an independent statistical test.

**Figure 2.**
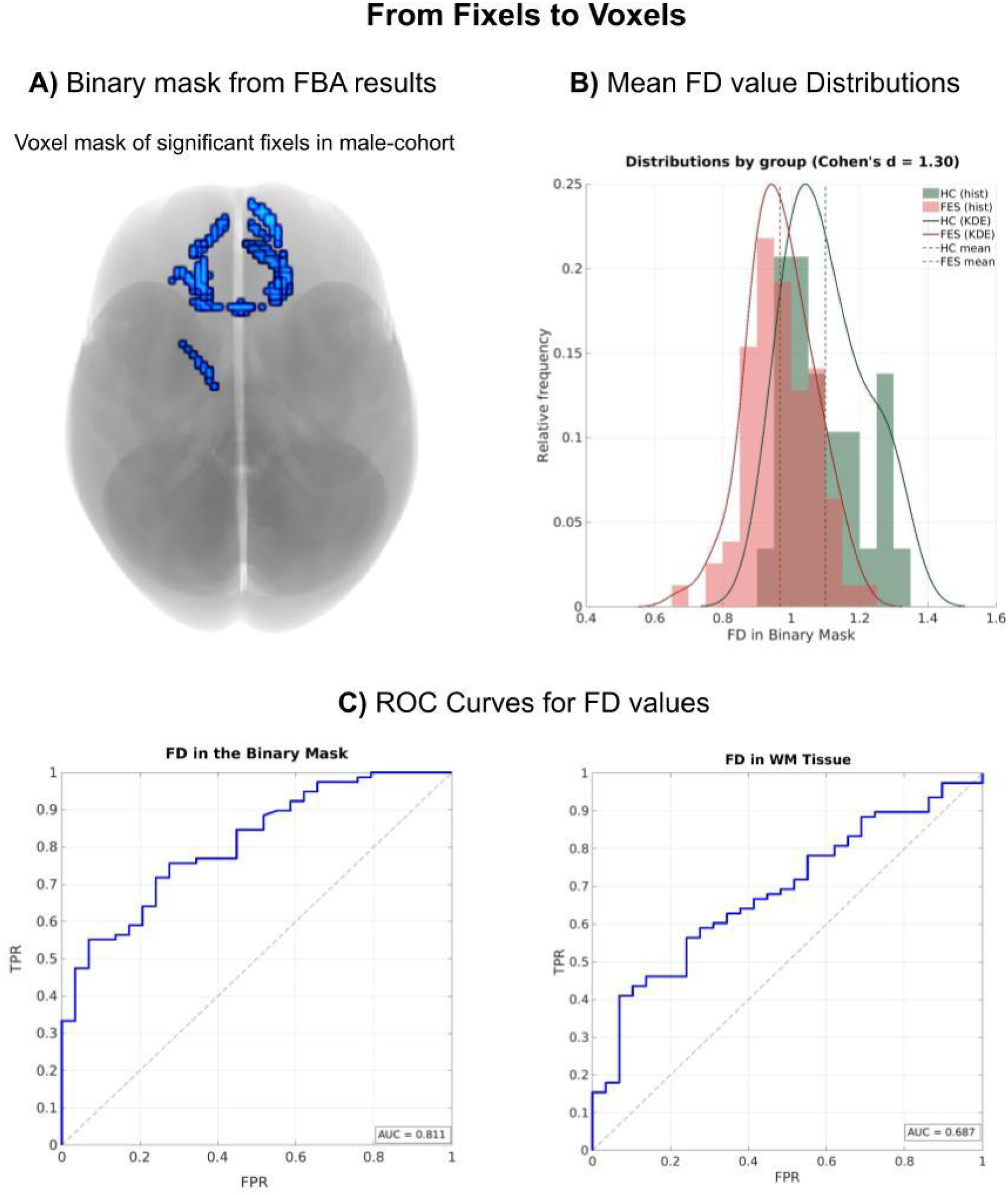
From Fixel to Voxel Scale. **(A)** Binary voxel mask derived from significant fixels in male-cohort in axial view. (B) Distributions of mean FD values per participant extracted from the binary mask, with FES shown in pink and HC in green. (C) ROC Curves computed for FD values. Left panel: In the binary mask in male-template space. AUC = 0.811. Right panel: Within WM tissue of the whole brain. AUC = 0.686. Both plots show TPR (sensitivity) vs. FPR. Abbreviations: FBA, Fixel-Based Analysis; FD, Fibre Density; HC, Healthy Controls; FES, First-Episode Schizophrenia; KDE, Kernel Density Estimation; ROC, Receiver Operating Characteristic; AUC, Area Under the Curve; WM, White Matter; TPR, True Positive Rate; FPR, False Positive Rate.

When calculating the ROC curve, we found that the FD from the binary mask defined after FBA in the male cohort successfully discriminates between male-FES and male-HC with AUC=0.811 (Figure 2C, left). To put the performance in context, we also calculated the AUC of FD values from the whole WM tissue, yielding AUC=0.686 (Figure 2C, right) and *d*=0.65. Note that this whole WM analysis does not rely on a prior region selection.

### 3.3 FBA on Female Cohort

FBA yielded no significant differences in the female cohort in any of the three fibre-specific metrics.

As these negative results might be putatively ascribed to the relatively low power of whole-brain fixel-based analysis, we further investigated whether specifically the WM area defined by the FBA outcome in the male cohort (Figure 1) may exhibit any substantial alterations in the female cohort when studied in isolation from the rest of the brain. To achieve this, we applied a non-linear 4D deformation field to warp the binary mask, defined in the male-template space, in order to map it to the female-template space. Once the binary mask was converted to the female-template space, we extracted the FD values from both groups (female-FES and female-HC) and made the appropriate comparison.

The sensitivity power analysis indicated that with 46 HC and 48 FES females, power would have been ∼100% to detect an effect of the magnitude observed in males (*d*=1.3, blue marker in Figure 3). Considering that the WM area defined in the male cohort may inflate the observed male effect size, power was also estimated for smaller effects: 87.6% for *d*=0.65 (green marker in Figure 3) and 66.9% for *d*=0.5 (red marker in Figure 3). With the current female sample size, 80% power corresponds to approximately *d*≈0.58 (orange dashed line in Figure 3). The observed effect size in females (*d*=0.12) is indicated by a purple diamond.

**Figure 3.**
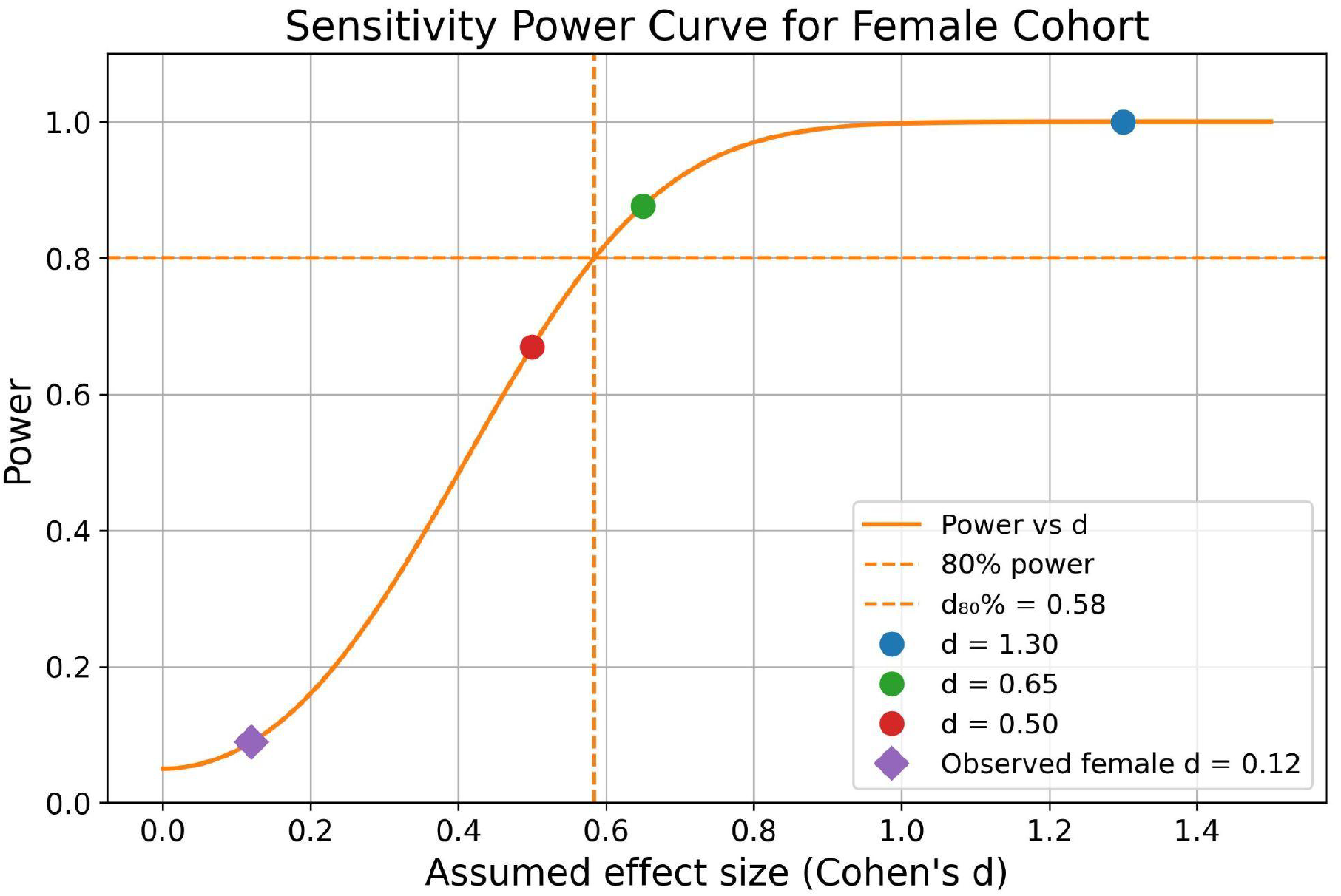
Sensitivity power curve for the female cohort. The curve shows statistical power for an independent two-sample t-test (two-sided α=0.05) across a range of assumed standardised effect sizes (Cohen’s d), given the female sample sizes (HC: n=46; FES: n=48). The dashed horizontal line indicates 80% power, and the dashed vertical line marks the effect size required to achieve 80% power (d_80%_ ≈0.58). Circles indicate the effect size observed in males (d=1.3) and more conservative assumptions (d=0.65 and d=0.5), while the purple diamond indicates the observed effect size in females (d=0.12) from the binary-mask-based comparison.

However, even at the voxel scale and in the average over the area defined by the binary mask, we detected no differences between the female groups in the FD metric (FD_HC_=0.89 ± 0.079; FD_FES_=0.88 ± 0.079; *p*=0.535, *d*=0.12). For comparison, the FD metric for the whole WM tissue in females yielded no significant differences and a small effect size: FD_HC_=0.64 ± 0.043; FD_FES_=0.63 ± 0.039; *p*=0.684, *d*=0.10.

## Discussion

In this study, we applied Fixel-Based Analysis (FBA) together with statistical methods to quantify white matter (WM) microstructure and examine group differences using a sex-stratified approach in individuals who had experienced a first episode of schizophrenia (FES). The sample was stratified by biological sex, and each subgroup was compared with its diagnostic counterpart, yielding four groups: male-FES, male-HC, female-FES, and female-HC.

First, we analysed demographic and MRI-derived variables to assess the characteristics and quality of the sample. We found no significant between-group differences in age, although a trend was observed in the male cohort. Years of education differed significantly between groups in the male cohort, whereas this effect was not observed in females. Differences in educational attainment between healthy controls and patients with schizophrenia have been consistently reported over the past decades, with lower educational attainment being associated with poorer health and well-being outcomes (23,25). In this context, years of education may partly reflect disorder-related characteristics rather than an independent confounding factor. Accordingly, matching or adjusting for years of education may introduce bias by favouring the selection of atypical subgroups, such as high-achieving patients or low-achieving controls. Therefore, years of education was not included as a covariate in the subsequent analyses.

In contrast, head motion was considered a potential confounder, as it is well established that motion can affect MRI acquisitions, not only by impairing image alignment but also by influencing the intensity of derived measurements (24). In diffusion MRI, motion during diffusion-encoding gradient pulses can lead to signal attenuation and, consequently, bias diffusion-derived metrics. To minimise these effects, we implemented two widely used strategies: first, DWIs were registered to a baseline image to correct for motion-related misalignment (26,27); and second, motion-related measures were included as nuisance regressors in the statistical analyses (24). The latter represents a conservative treatment of motion-related artefacts, reducing the likelihood that the observed effects were driven by residual motion differences.

Clinically, our FES sample did not show significant differences between males and females in age at onset of symptoms or in PANSS positive, negative, or general psychopathology scores. As outlined in the Introduction, previous studies have reported an earlier age at onset in males with schizophrenia, together with sex-related differences in symptom profiles. However, such differences appear less consistent in first-episode samples. For instance, Zhang and colleagues (28) reported an earlier onset in males with first-episode schizophrenia but no sex differences in symptom severity, while Cocchi et al. (29) reported no significant diagnosis-by-sex interaction in symptom severity.

Our findings revealed WM microstructural abnormalities in males with FES relative to male-HC. More specifically, the male-FES group exhibited reduced fibre density (FD), a proxy of the intra-axonal volume of a fibre population, suggesting lower apparent axonal content within the affected fibre bundles. These reductions were predominantly located in bilateral frontal WM, adjacent to cortical regions including the superior frontal gyrus, rostral anterior cingulate, and orbitofrontal cortex. This pattern is noteworthy, as frontal cortical regions have been consistently implicated in schizophrenia and associated with early symptom expression, structural abnormalities, and impairments in social and cognitive functioning (30,31).

The present findings are in agreement with previous literature indicating that schizophrenia is characterised by alterations in frontal and prefrontal WM, as well as in grey matter structures (32,33). Quantitative structural MRI studies have further reported grey matter volume loss in thalamocortical connections and in the prefrontal cortex (14). Although, it is not fully understood how schizophrenia would be associated to damage or decreased WM density (34), histological evidence suggests that such grey matter reductions are accompanied by decreases in dendritic and synaptic density, which may contribute to aberrant neural communication and are consistent with the disconnection hypothesis of schizophrenia (35). In addition, postmortem findings support the possibility of primary structural abnormalities in the prefrontal cortex (36–39).

By contrast, the female cohort did not show significant group differences in any of the three FBA-derived metrics. Moreover, even when the analysis was restricted to the FD-altered region identified in males, no significant FD differences were observed between female-FES and female-HC. The sensitivity power analysis further indicated that the available female sample was sufficiently powered to detect an effect of a magnitude comparable to that observed in males, making a large male-like effect in females less likely. Taken together, these findings are consistent with the possibility of sex-related differences in the expression or timing of WM alterations, in line with evidence suggesting an earlier clinical onset in males (3). Interestingly, these FD differences are unlikely to be driven by sex differences in the clinical characteristics examined here, as males and females with FES did not differ significantly in age at onset or PANSS scores as reported above.

From a technical perspective, several points should also be emphasised. The FBA presents two fundamental advantages over conventional voxel-wise diffusion methods: sensitivity to microstructure-specific properties that is less confounded by local fibre geometry, and fibre-specific analysis and interpretation, allowing effects associated with individual fibre populations within a voxel to be identified (40). By contrast, conventional diffusion-derived scalar metrics can lead to misleading or non-intuitive biophysical interpretations (41,42), particularly because they are not specific to axonal properties. Finally, it should be emphasised that the voxel-scale FD comparison was not independent of the primary FBA, as the mask used for FD extraction was defined from the thresholded significant fixels. Accordingly, this analysis should be interpreted as a descriptive voxel-scale representation of the fixel-level findings rather than as an independent replication of the group effect.

To the best of our knowledge, no recent studies have applied an updated sex-stratified approach of this kind in this context. We acknowledge that this methodological decision may be open to criticism; however, the underrepresentation of women in neuroscience (43–45) and clinical research (46) remains a persistent limitation that may contribute to inaccurate diagnoses and suboptimal treatment strategies. In this regard, stratifying the cohort by sex enables a more refined anatomical characterisation of each subgroup and may help establish a foundation for future investigations.

An important consideration when interpreting the sex-stratified findings is that evidence for a significant diagnostic effect in one sex and the absence of a significant effect in the other do not, by themselves, demonstrate that the diagnostic effect differs significantly between sexes. Accordingly, our findings should be interpreted as evidence of reduced frontal FD in males with FES, whereas no corresponding FD alteration was detected in females. This distinction limits conclusions regarding a formally demonstrated sex-specific effect, while still highlighting a differential pattern emerging from the sex-stratified analyses.

Finally, replication in larger and adequately powered samples will be necessary to establish the robustness and specificity of the present findings. Our results are consistent, in males, with patterns of WM alteration previously reported in schizophrenia using other methodological approaches, whereas a comparable pattern was not detected in the female cohort. This raises the possibility that effects reported in mixed-sex samples may, in some instances, be influenced more strongly by male participants, although this hypothesis requires direct testing in independent cohorts. Despite the sensitivity analysis indicating adequate power to detect a male-sized effect in the female sample, larger female cohorts will be important for identifying potentially subtler alterations and for determining whether the apparent divergence between sexes is reproducible. In addition, larger samples will be needed to disentangle potential confounding factors related to biological sex, illness stage, and treatment exposure, and to explicitly address the role of gender in addition to biological sex. A clearer understanding of how early white matter alterations unfold, and whether these trajectories differ according to sex and gender, may ultimately inform the development of more targeted and stage-sensitive interventions.

## Acknowledgments

The authors would like to sincerely thank all subjects for their time and effort.

## Ethical statement

The authors confirm that all experiments involving human participants were performed following relevant guidelines and regulations. The study was conducted according to the guidelines of the Declaration of Helsinki and approved by the Ethics Committee of the Prague Psychiatric Centre (protocol code 69/11, approved on 29 June 2011). Written informed consent was obtained from all subjects involved in the study.

## References

1. Giordano GM, Bucci P, Mucci A, Pezzella P, Galderisi S (2021): Gender Differences in Clinical and Psychosocial Features Among Persons With Schizophrenia: A Mini Review. Front Psychiatry 12. 10.3389/fpsyt.2021.789179

2. Li X, Zhou W, Yi Z (2022): A glimpse of gender differences in schizophrenia. Gen Psychiatry 35: e100823.

3. McGrath J, Saha S, Chant D, Welham J (2008): Schizophrenia: A Concise Overview of Incidence, Prevalence, and Mortality. Epidemiol Rev 30: 67–76.

4. Aleman A, Kahn RS, Selten J-P (2003): Sex Differences in the Risk of Schizophrenia: Evidence From Meta-analysis. Arch Gen Psychiatry 60: 565–571.

5. Friston K (2002): Dysfunctional connectivity in schizophrenia. World Psychiatry 1: 66–71.

6. Kelly S, Jahanshad N, Zalesky A, Kochunov P, Agartz I, Alloza C, et al. (2018): Widespread white matter microstructural differences in schizophrenia across 4322 individuals: results from the ENIGMA Schizophrenia DTI Working Group. Mol Psychiatry 23: 1261–1269.

7. Konrad A, Winterer G (2008): Disturbed Structural Connectivity in Schizophrenia—Primary Factor in Pathology or Epiphenomenon? Schizophr Bull 34: 72–92.

8. Assaf Y, Johansen-Berg H, Thiebaut de Schotten M (2019): The role of diffusion MRI in neuroscience. NMR Biomed 32: e3762.

9. Lerner A, Mogensen MA, Kim PE, Shiroishi MS, Hwang DH, Law M (2014): Clinical Applications of Diffusion Tensor Imaging. World Neurosurg 82: 96–109.

10. Figley CR, Uddin MN, Wong K, Kornelsen J, Puig J, Figley TD (2022): Potential Pitfalls of Using Fractional Anisotropy, Axial Diffusivity, and Radial Diffusivity as Biomarkers of Cerebral White Matter Microstructure. Front Neurosci 15. 10.3389/fnins.2021.799576

11. Schilling KG, Tax CMW, Rheault F, Landman BA, Anderson AW, Descoteaux M, Petit L (2022): Prevalence of white matter pathways coming into a single white matter voxel orientation: The bottleneck issue in tractography. Hum Brain Mapp 43: 1196–1213.

12. Raffelt DA, Smith RE, Ridgway GR, Tournier J-D, Vaughan DN, Rose S, et al. (2015): Connectivity-based fixel enhancement: Whole-brain statistical analysis of diffusion MRI measures in the presence of crossing fibres. NeuroImage 117: 40–55.

13. Agartz I, Andersson JLR, Skare S (2001): Abnormal brain white matter in schizophrenia: a diffusion tensor imaging study. NeuroReport 12: 2251.

14. Dabiri M, Dehghani Firouzabadi F, Yang K, Barker PB, Lee RR, Yousem DM (2022): Neuroimaging in schizophrenia: A review article. Front Neurosci 16. 10.3389/fnins.2022.1042814

15. Melicher T, Horacek J, Hlinka J, Spaniel F, Tintera J, Ibrahim I, et al. (2015): White matter changes in first episode psychosis and their relation to the size of sample studied: A DTI study. Schizophr Res 162: 22–28.

16. Mikolas P, Hlinka J, Skoch A, Pitra Z, Frodl T, Spaniel F, Hajek T (2018): Machine learning classification of first-episode schizophrenia spectrum disorders and controls using whole brain white matter fractional anisotropy. BMC Psychiatry 18: 97.

17. Lecrubier Y, Sheehan D, Weiller E, Amorim P, Bonora I, Harnett Sheehan K, et al. (1997): The Mini International Neuropsychiatric Interview (MINI). A short diagnostic structured interview: reliability and validity according to the CIDI. Eur Psychiatry 12: 224–231.

18. Škoch A, Rehák Bučková B, Mareš J, Tintěra J, Sanda P, Jajcay L, et al. (2022): Human brain structural connectivity matrices–ready for modelling. Sci Data 9: 486.

19. Jeurissen B, Tournier J-D, Dhollander T, Connelly A, Sijbers J (2014): Multi-tissue constrained spherical deconvolution for improved analysis of multi-shell diffusion MRI data. NeuroImage 103: 411–426.

20. Sánchez SM, Knösche TR, Fürstová P, Škoch A, Španiel F, Schmidt H, Hlinka J (2025): Characterising structural brain connectivity of patients with first episode of psychosis. bioRxiv 2025–05.

21. Raffelt D, Dhollander T, Tournier J-D, Tabbara R, Smith R, Pierre E, Connelly A (2017): Bias Field Correction and Intensity Normalisation for Quantitative Analysis of Apparent Fibre Density.

22. Bastiani M, Cottaar M, Fitzgibbon SP, Suri S, Alfaro-Almagro F, Sotiropoulos SN, et al. (2019): Automated quality control for within and between studies diffusion MRI data using a non-parametric framework for movement and distortion correction. NeuroImage 184: 801–812.

23. Crossley NA, Alliende LM, Czepielewski LS, Aceituno D, Castañeda CP, Diaz C, et al. (2022): The enduring gap in educational attainment in schizophrenia according to the past 50 years of published research: a systematic review and meta-analysis. Lancet Psychiatry 9: 565–573.

24. Yendiki A, Koldewyn K, Kakunoori S, Kanwisher N, Fischl B (2014): Spurious group differences due to head motion in a diffusion MRI study. NeuroImage 88: 79–90.

25. Hakulinen C, McGrath JJ, Timmerman A, Skipper N, Mortensen PB, Pedersen CB, Agerbo E (2019): The association between early-onset schizophrenia with employment, income, education, and cohabitation status: nationwide study with 35 years of follow-up. Soc Psychiatry Psychiatr Epidemiol 54: 1343–1351.

26. Andersson JLR, Skare S (2002): A Model-Based Method for Retrospective Correction of Geometric Distortions in Diffusion-Weighted EPI. NeuroImage 16: 177–199.

27. Rohde G k., Barnett A s., Basser P j., Marenco S, Pierpaoli C (2004): Comprehensive approach for correction of motion and distortion in diffusion-weighted MRI. Magn Reson Med 51: 103–114.

28. Zhang X, Chen D, Xiu M, Yang F, Haile C, Kosten T, Kosten T (2012, July): Gender Differences in Never-Medicated First-Episode Schizophrenia and Medicated Chronic Schizophrenia Patients. 10.4088/JCP.11m07422

29. Cocchi A, Lora A, Meneghelli A, La Greca E, Pisano A, Cascio MT, Preti A (2014): Sex differences in first-episode psychosis and in people at ultra-high risk. Psychiatry Res 215: 314–322.

30. Jung H-Y, Jung S, Bang M, Choi TK, Park CI, Lee S-H (2022): White matter correlates of impulsivity in frontal lobe and their associations with treatment response in first-episode schizophrenia. Neurosci Lett 767: 136309.

31. Leucht S, Siafis S, McGrath JJ, McGorry P, Howes OD, Tamminga C, et al. (2025): Schizophrenia. Nat Rev Dis Primer 11: 83.

32. Breier A, Buchanan RW, Elkashef A, Munson RC, Kirkpatrick B, Gellad F (1992): Brain morphology and schizophrenia. A magnetic resonance imaging study of limbic, prefrontal cortex, and caudate structures. Arch Gen Psychiatry 49: 921–926.

33. Buchanan RW, Vladar K, Barta PE, Pearlson G (1998): Structural Evaluation of the Prefrontal Cortex in Schizophrenia. Am J Psychiatry. Retrieved April 13, 2026, from https://psychiatryonline.org/doi/10.1176/ajp.155.8.1049

34. Grazioplene RG, Bearden CE, Subotnik KL, Ventura J, Haut K, Nuechterlein KH, Cannon TD (2018): Connectivity-enhanced diffusion analysis reveals white matter density disruptions in first episode and chronic schizophrenia. NeuroImage Clin 18: 608–616.

35. Glantz LA, Lewis DA (2000): Decreased Dendritic Spine Density on Prefrontal Cortical Pyramidal Neurons in Schizophrenia. Arch Gen Psychiatry 57: 65–73.

36. Akbarian S, Bunney WE, Potkin SG, Wigal SB, Hagman JO, Sandman CA, Jones EG (1993): Altered distribution of nicotinamide-adenine dinucleotide phosphate-diaphorase cells in frontal lobe of schizophrenics implies disturbances of cortical development. Arch Gen Psychiatry 50: 169–177.

37. Benes FM, McSparren J, Bird ED, SanGiovanni JP, Vincent SL (1991): Deficits in small interneurons in prefrontal and cingulate cortices of schizophrenic and schizoaffective patients. Arch Gen Psychiatry 48: 996–1001.

38. Daviss SR, Lewis DA (1995): Local circuit neurons of the prefrontal cortex in schizophrenia: selective increase in the density of calbindin-immunoreactive neurons. Psychiatry Res 59: 81–96.

39. Selemon LD, Rajkowska G, Goldman-Rakic PS (1995): Abnormally high neuronal density in the schizophrenic cortex. A morphometric analysis of prefrontal area 9 and occipital area 17. Arch Gen Psychiatry 52: 805–818; discussion 819-820.

40. Dhollander T, Clemente A, Singh M, Boonstra F, Civier O, Duque JD, et al. (2021): Fixel-based Analysis of Diffusion MRI: Methods, Applications, Challenges and Opportunities. NeuroImage 241: 118417.

41. Beaulieu C (2014): Chapter 8 - The Biological Basis of Diffusion Anisotropy. In: Johansen-Berg H, Behrens TEJ, editors. Diffusion MRI (Second Edition). San Diego: Academic Press, pp 155–183.

42. Jones DK, Christiansen KF, Chapman RJ, Aggleton JP (2013): Distinct subdivisions of the cingulum bundle revealed by diffusion MRI fibre tracking: Implications for neuropsychological investigations. Neuropsychologia 51: 67–78.

43. Geller SE, Koch AR, Roesch P, Filut A, Hallgren E, Carnes M (2018): The More Things Change, the More They Stay the Same: A Study to Evaluate Compliance With Inclusion and Assessment of Women and Minorities in Randomized Controlled Trials. Acad Med 93: 630–635.

44. Pederson SL, Lindstrom R, Powe PM, Louie K, Escobar-Viera C (2022): Lack of Representation in Psychiatric Research: A Data-Driven Example From Scientific Articles Published in 2019 and 2020 in the American Journal of Psychiatry. Am J Psychiatry. Retrieved April 13, 2026, from https://psychiatryonline.org/doi/10.1176/appi.ajp.21070758

45. Woodall A, Morgan C, Sloan C, Howard L (2010): Barriers to participation in mental health research: are there specific gender, ethnicity and age related barriers? BMC Psychiatry 10: 103.

46. Daitch V, Turjeman A, Poran I, Tau N, Ayalon-Dangur I, Nashashibi J, et al. (2022): Underrepresentation of women in randomized controlled trials: a systematic review and meta-analysis. Trials 23: 1038.

